# 12-plex DiLeu enables robust quantification of the feeding neuropeptidome

**DOI:** 10.64898/2026.08.05.743115

**Authors:** Lauren Fields, Peng-Kai Liu, Vu Ngoc Huong Tran, Thao Duong, Angel E. Ibarra, Paul Corsetti, Tong Gao, Kendra G. Selby, Zicong Wang, Yajing Lu, Haiyan Lu, Lingjun Li

## Abstract

Understanding the feeding-induced neuropeptidome cascade requires analytical strategies capable of quantifying low-abundance, highly modified peptides across multiple tissues and time points. Herein, we apply 12-plex *N,N*-dimethyl leucine (DiLeu) isobaric labeling to perform the first multiplexed, tissue-wide, temporal quantitation of the *Cancer borealis* feeding neuropeptidome. This approach enabled sensitive measurement of neuropeptides across five tissues over six timepoints, revealing distinct regulatory patterns. The pericardial organ (PO) showed rapid early upregulation followed by suppression aligned with foregut emptying, whereas the thoracic ganglion (TG) displayed inverse and strongly condition-dependent responses, indicating previously unrecognized neuromodulatory roles. Single-residue variants and post-translational modifications, including pyro-Glu formation and amidation, produced markedly different temporal profiles, underscoring the functional specificity of closely related isoforms. We further identify differential regulation of proctolin and its amidated form, suggesting modified variants may contribute uniquely to feeding physiology. Collectively, these results establish multiplexed DiLeu labeling as a powerful platform for quantitative neuropeptidomics and reveal new dimensions of peptide-mediated feeding regulation.

## Introduction

Feeding processes are a common thread underlying many conditions and diseases impacting global health, including diabetes, anorexia, and bulimia. Despite this, the mechanistic underpinnings of feeding remains largely uncharacterized. However, specific neuronal circuits, such as those governing the gastric mill (*i*.*e*. chewing) and pyloric rhythm (*i*.*e*. filtering of food) have been extensively studied in the Jonah crab, *Cancer borealis*, providing a well-defined model system for understanding motor pattern generation and modulation.^1^ We now understand neuropeptides heavily modulate these processes, though many co-modulatory and circuit-level interactions remain poorly defined.

In recent decades, mass spectrometry-based neuropeptidomics has become a powerful strategy for interrogating neuropeptide signaling.^2^ For increased throughput and reliability, isobaric labeling has proven to be an appealing strategy, addressing common quantification pitfalls such as retention time shifts and quantification pitfalls such as retention time shifts and run-to-run variation. Despite this, isobaric labeling strategies have only rarely been applied to neuropeptides.^2-4^ Most routinely, isotopic chemical labeling has been used, but these approaches typically offer limited multiplexing capability.^5, 6^ As a cost-effective alternative to commercial isobaric labels such as TMT, DiLeu offers a simple synthetic scheme that enables robust isobaric labeling in up to 21-plex labeling regimes.^7, 8^

Multifaceted strategies have been employed to evaluate feeding-related signaling within *Cancer borealis* and other crustacea, including mass spectrometry imaging,^1, 3^ isotopic labeling, capillary electrophoresis,^3^ data-independent acquisition mass spectrometry,^4^ and more. These innovative methods have revealed that the hormonal response is dynamically regulated by neuropeptides throughout the chewing and digestion processes.^5^ This renders *Cancer borealis* an excellent system for dissecting key neuropeptidergic players involved in feeding.

Here, we employ 12-plex DiLeu isobaric labeling to perform relative quantitation of neuropeptides across multiple tissues and time points spanning the feeding cycle. This approach allows us to reveal neuromodulation imparted throughout the feeding domain, with a particular emphasis on the pericardial organ (PO) and thoracic ganglion (TG), which emerge as major sites of dynamic neuropeptide regulation. We further interrogate the contribution of closely related isoforms and post-translationally modified variants to feeding-associated signaling, and we uncover evidence that coordinated neuropeptide responses extend beyond the canonical 24-hour post-feeding window.

### Experimental Section

#### Chemicals and Reagents

ACS grade chemicals, including 4-(4,6-Dimethoxy-1,3,5-triazin-2-yl)-4-methylmorpholinium tetrafluoroborate (DMTMM BF4), dimethylformamide (DMF), 4-methylmorpholin (NMM), 1.0 M triethylammonium bicarbonate buffer (TEAB), methanol (MeOH), hydroxylamine (NH2OH), ammonium hydroxide (NH4OH), formic acid (FA), trifluoroacetic acid (TFA), water (H2O), and acetonitrile (ACN) were purchased from Sigma Aldrich (St. Louis, MO). Optima-grade solvents for MS analysis were purchased from Fisher Scientific (Pittsburgh, PA). Acidified methanol was prepared with 90/9/1 of H2O/MeOH/acetic acid.

#### Animal Experiments

Male Jonah crabs (*Cancer borealis*) were purchased from a local vendor (Global Markets, Madison, WI), ensuring similar size and health for all organisms. Crabs were stored in an artificial seawater environment, utilizing a salt mixture (Instant Ocean, Blacksburg, VA) at a concentration of 35 parts per thousand salinity. Additionally, the environment was maintained at 13-16 °C with 8-10 ppm circulating O2. Crabs were equilibrated in this environment for 2 weeks without feeding to ensure stomachs were empty. For feeding, crabs were provided 4 g thawed tilapia, consistent with the quantity required for satiety.^4^ Crabs were permitted to eat for 30 min, and then were prepared for dissection following the intended time (*e*.*g*. 0 h, 30 min, 1 h, 4 h, 6 h, 24 h). For each fed crab, an unfed crab was also sacrificed at the same timepoint. Crabs were anesthetized on packed ice for 30 min prior to dissection.

#### Tissue Extraction

Sinus gland (SG), pericardial organ (PO), thoracic ganglion (TG), brain, and stomatogastric nervous system (STNS) tissue were collected from fed and unfed crabs. Immediately, tissues were heat-stabilized via Denator, and flash frozen to -80 °C. Neuropeptides were extracted in acidified methanol (see *Chemicals* subsection) via probe sonication, with like tissue pooled, and promptly dried. Subsequently, neuropeptides were desalted with C18 ZipTips (Agilent, Santa Clara, CA), releasing in a gradient of 25%, 50%, and 75% ACN. Each fraction was pooled and dried down prior to labeling.

#### DiLeu Labeling

12-plex DiLeu tags were prepared, with the synthesis described elsewhere.^12^ Of the 12 DiLeu channels, 6 were reserved for control tissue, and 6 were for fed tissue, with three tissue pooled per channel. For each channel, 0.5 mg DiLeu tags were dried to ensure no H2O presence. An activation solution was prepared with 12.9 mg DMTMM BF4, 412.5 μL DMF, and 4.3 μL NMM. To each tag, 25 μL activation solution was added, and activation was conducted by vortexing in darkness for 50 min. Peptide samples for labeling were reconstituted in 5 μL 0.5 M TEAB buffer. For labeling, activated tags were added to reconstituted peptide samples, and labeling was carried out by vortexing in darkness for 2 h. Reactions were then promptly quenched with NH2OH to a final NH2OH concentration of 0.25%. All 12 samples were then combined and dried down. Subsequently, strong cation exchange (SCX) cleanup was conducted with OMIX SCX Tips (Agilent, Santa Clara, CA). The labeled sample was reconstituted in 100 μL 0.1% TFA to a pH less than 4.0. Cleanup was performed according to manufacturer instructions, eluting in 100 μL 5% NH4OH in 30% MeOH. The eluent was dried down, after which the sample then underwent C18 desalting as before.

#### LC-MS/MS Analysis

Sample analysis was performed using a Dionex UltiMate 3000 nanoLC system coupled to a Fusion Lumos Tribrid mass spectrometer (Thermo Fisher Scientific, San Jose, CA). Mobile phases A and B were comprised of 0.1% FA in H2O and 0.1% FA in ACN, respectively. Online chromatography was conducted with a homemade C18 column at a flow rate of 0.25 µL/min. A 138 min gradient started at 4% B and ramped to 50% B in 112 min. Mass spectrometry was conducted in positive mode, with the following parameters: MS1 Orbitrap resolution, 60K; scan range, 200-2000 m/z, RF lens, 30%; maximum injection time, 250 ms; intensity threshold, 1000; dynamic exclusion duration, 45 s. The top 20 precursors were selected for MS^2^, as well as the following parameters for MS^2^: isolation window, 2 *m/z*; HCD collision energy, 30%; Orbitrap resolution, 60K; fixed first mass, 100 *m/z*; maximum injection time, 118 ms. All samples were acquired in technical triplicate.

#### Data analysis

Thermo .RAW files were converted to .mzML and .MS2 formats via MSconvert^13^ and RawConverter,^14^ respectively, using default settings. Analyses were conducted in EndoGenius 1.14,^15^ available freely at (https://github.com/lingjunli-research/EndoGenius), as described previously.^15-17^ Specifically, precursor and fragment error tolerance were set to 30 ppm and 0.05 Da, respectively. 12-plex DiLeu was indicated as a fixed modification, with C-terminal amidation, methionine oxidation, and pyro-Glu of Q and E as variable modifications. A maximum of 5 modifications were permitted per peptide, with DiLeu labeling of the N-termini and lysine considered in this threshold. An EndoGenius score threshold of 750 was selected.

#### Statistical analysis

Reporter ion intensities were normalized as described previously.^18^ Isotopic corrections were applied using the matrix provided in **Table S1**. Statistics were applied via BioRender statistics package, performing a 2-way ANOVA with Tukey multiple comparisons test.

## Results and Discussion

Multiplexing via isobaric labeling offers tremendous opportunity to simultaneously probe complex biological questions, such as the neuroendocrine response to feeding. Historically, the extent of multiplexing for neuropeptides has largely been applied with isotopic labeling strategies,^1, 3^ which inherently are limited in the number of available channels. In part, the limited adoption of more advanced isobaric quantitation methods for neuropeptides reflects the relatively large peptide amounts typically required. For example, standard isobaric labeling workflows are often designed for scenarios in which at least 50 μg of starting material is available, an amount that is infeasible for crustacean samples.^18^ However, recent work has since made headway in applying this innovative method to limited samples, including single cells.^19-21^ Thus, we hypothesized that pooling three tissue bioreplicates per channel would provide sufficient aggregate peptide load to overcome the low abundance of neuropeptides.

To overcome challenges in abundance, three crabs were pooled per channel (**Fig. 1**). Largely, the protocol for labeling was consistent with those used for proteins. However, the downstream data analysis was tailored specifically for neuropeptidomics. We have shown in several instances that EndoGenius, a software platform optimized for analysis of endogenous peptides, substantially improves neuropeptide detection compared to other commonly employed methods.^16, 17^ Recently, we reported the expansion of EndoGenius to include isobaric labeling workflows.^17^ One reason for the improved results with EndoGenius is the application of a custom EndoGenius score, a weighted value determined by hyperscore, motif scores, series of b- and y-ions, and additional spectral features.

**Figure 1.**
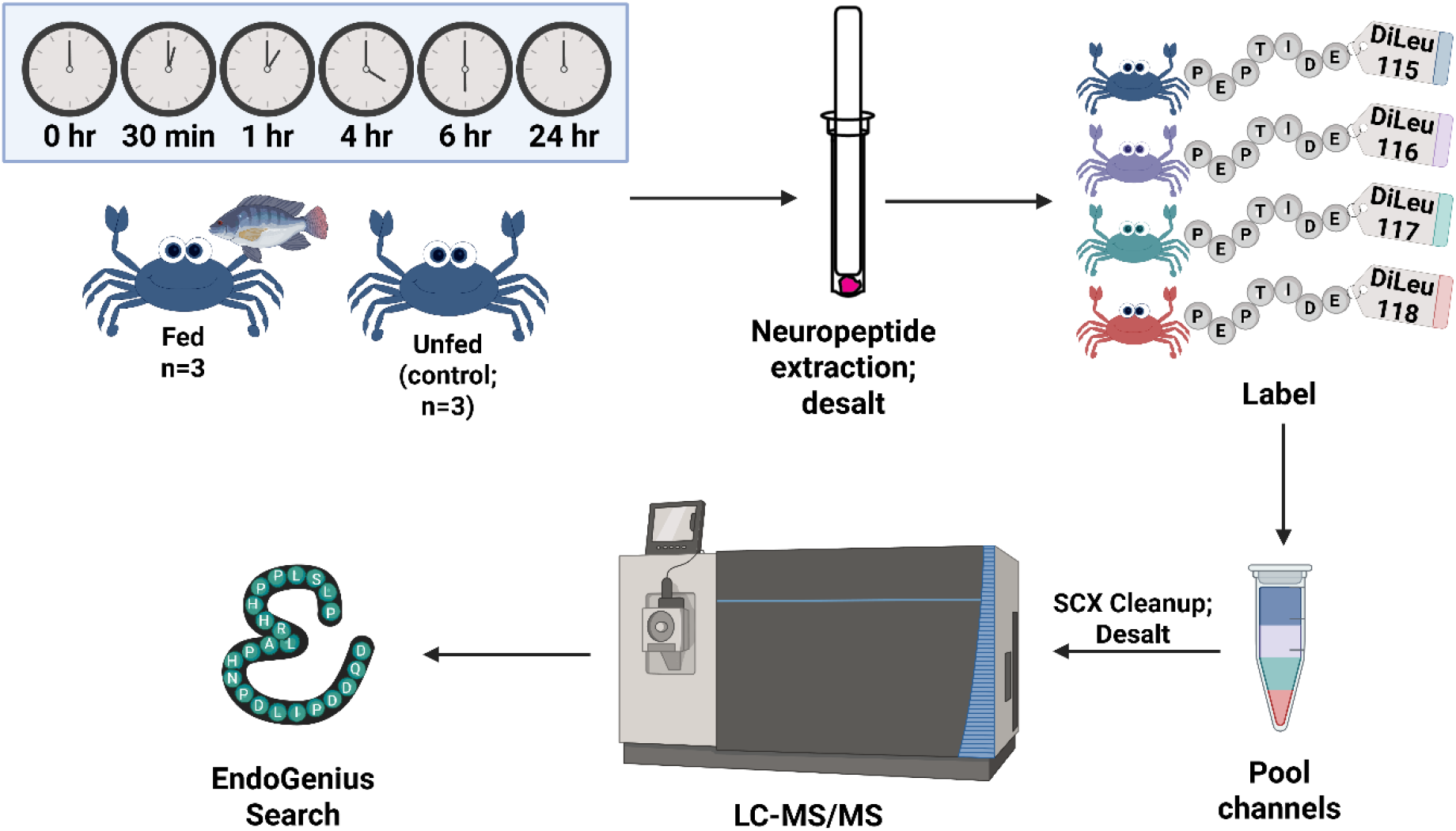
General workflow for labeling of *C. borealis* tissue throughout the feeding process. Of the 12 DiLeu channels, six were reserved for control tissue, and six were for fed tissue, with three tissues pooled per channel. (*i.e*., 18 tissues per feeding condition, 36 pooled total). Of the six channels per feeding condition, each represented a unique time point. Tissue extract was labeled, cleaned up, analyzed via LC-MS/MS, and evaluated through EndoGenius.

Given that neuropeptides are low in abundance and traditionally only a few hundred identifications are anticipated, a false-discovery rate (FDR) determined from decoy-to-target ratios can be highly variable, because the relatively small number of identifications amplifies the influence of decoy matches and can deviate from the expected FDR distribution. In lieu of FDR, we calibrate an EndoGenius score and employ a numeric threshold for identification, a strategy employed elsewhere.^22^ For instance, in applications for label-free analysis, a *log*(EndoGenius score) threshold of 1000 has been defined as a reliable value producing IDs below or equivalent to 5% FDR, the commonly used threshold for neuropeptidomics.^23^ In the work described herein, we recalibrated the EndoGenius score for application to DiLeu labeling. To do this, we pooled labeled SGs in equivalent ratios across all channels, as well as in a ratio of 1:2:4:6:10:12:12:10:6:4:2:1. Under these controlled conditions, where biological variance is minimized, we concluded that an EndoGenius score of 750 (*log*(EndoGenius score) ≈ 2.875) was appropriate for this application (**Fig. S1**).

Further, we assessed the fragmentation of neuropeptides, inclusive of the isobaric labels (**Fig 2**), before profiling and quantifying neuropeptides across the gamut of feeding times. We evaluated time points post-feeding at 0 h, 30 min, 1 h, 4 h, 6 h, and 24 h. In particular it is anticipated that the emptying of the foregut is about 6 h.^5^

**Figure 2.**
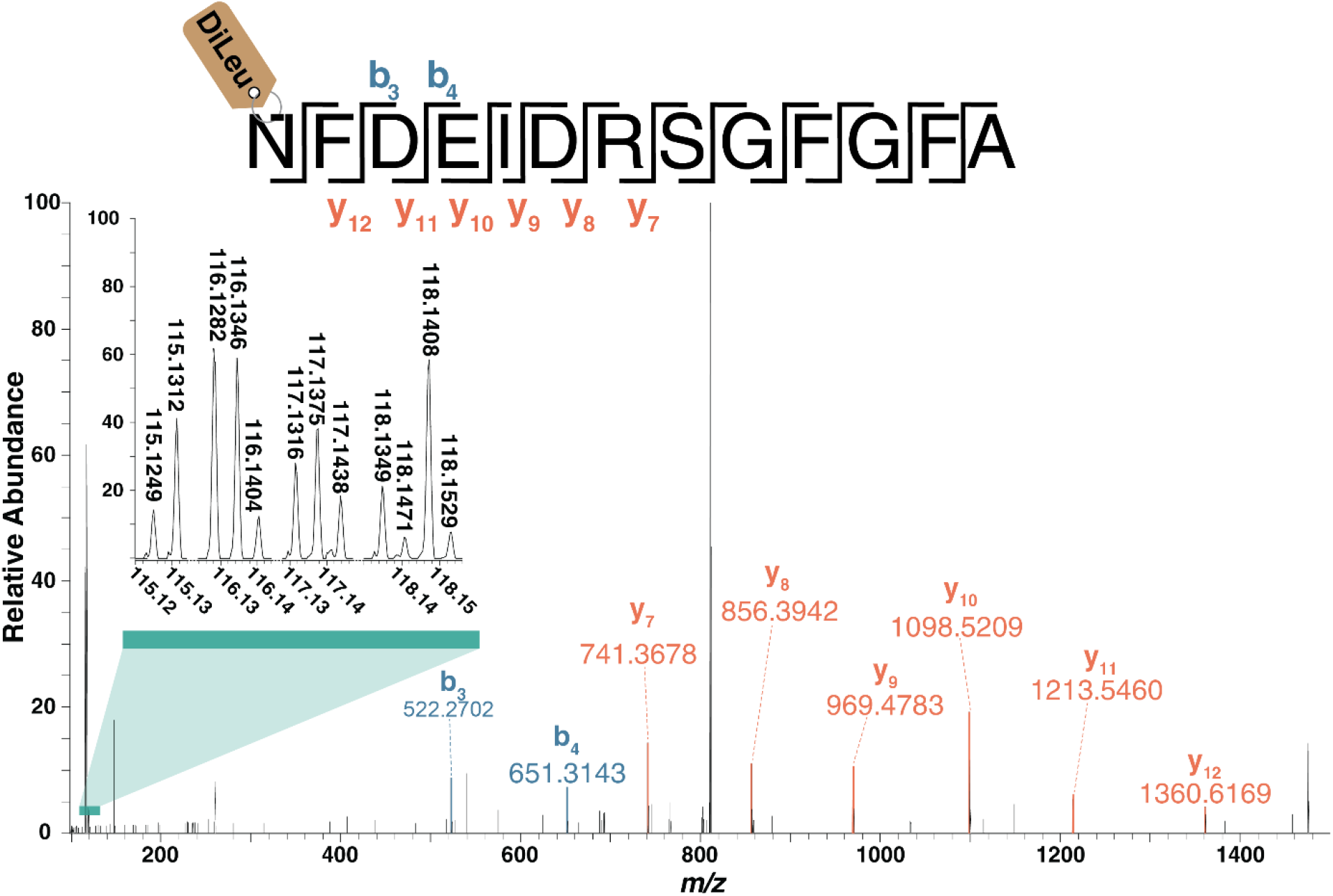
Representative annotated MS^2^ spectrum of labeled orcokinin neuropeptide NFDEIDRSGFGFA, including the entire set of reporter ions from 12-plex DiLeu. This peptide was identified in all conditions within the sinus gland (SG) tissues analyzed.

Upon evaluating the ratio of fed to unfed intensity for peptides across all tissue and time points, trends began to emerge (**Supplementary File 1**). The first notable observation came from NFLRF (**Fig. 3 A**). This neuropeptide, belonging to the RFamide family, albeit not observed in its amidated form, exhibited dramatic decrease in intensity within the PO from 1 to 4 h, as well as a significant increase in the TG within the first 30 minutes following feeding, and subsequent decrease by 6 h. RFamide neuropeptides, such as the one we observed, are strongly associated with feeding and among the most frequently observed neuropeptides across the animal kingdom.^2^ Part of the reason for this broad relevance is that it is conserved and homologous to neuropeptide Y in vertebrates.^2^ Previous work has extensively profiled several isoforms of NFLRFamide (*e.g*. NRNFLRFamide,^1, 24-33^ GNRNFLRFamide^1, 24, 25, 27, 31, 34^) within the blue crab, *Callinectes sapidus*, via mass spectrometry imaging, revealing a differential distribution of two peptides. This is representative of a common finding within neuropeptidomics, wherein peptides are incredibly homologous, and thus, it is difficult to determine if a peptide is merely a fragment, or if it is bioactive.^35^ As a result, peptide profiles typically encompass fragments, putative neuropeptides, validated neuropeptides, predicted peptides, and prohormone-related peptides.^36^

**Figure 3.**
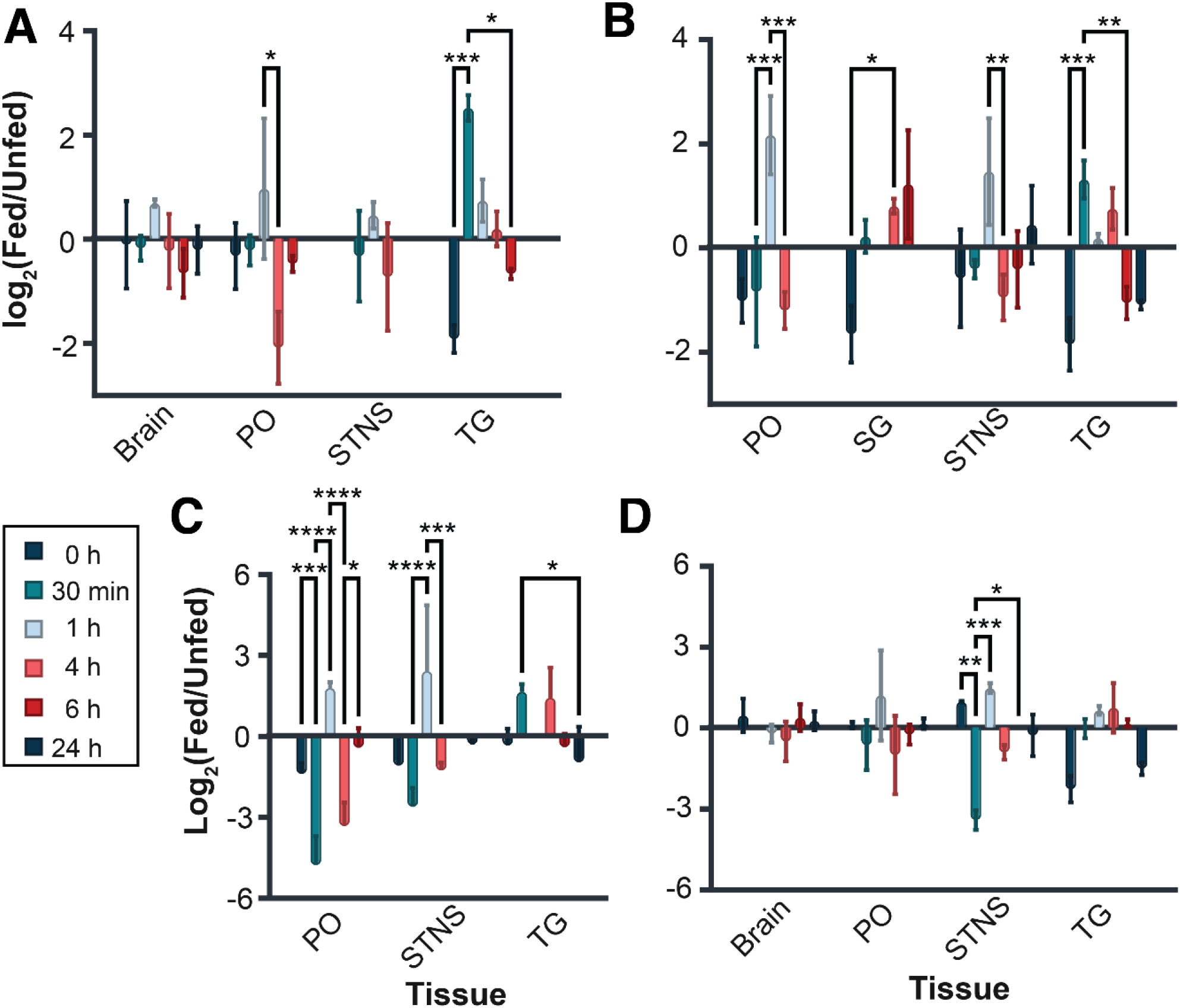
Ratio of fed to unfed peptide abundance across *C. borealis* tissue with respect to time for peptides **A)** NFLRF, **B)** SPYAFGLamide, **C)** KMYDFGLamide, and **D)** GIYGFGL within the brain, paired pericardial organs (PO), stomatogastric nervous system (STNS) and thoracic ganglion (TG). Bars represent the average of three technical replicates ± standard deviation. Significance was evaluated via two-way ANOVA with Bonferroni’s multiple comparison test. Tissues: pericardial organ (PO), sinus gland (SG), stomatogastric nervous system (STNS), thoracic ganglion (TG).

In another study, a series of these NFLRFamides were observed in the American Lobster, *Homarus americanus*.^35^ Given the dynamic nature of this peptide, we analyzed the other peptides containing the NFLRF subsequence in the current study. Within the PO, numerous significant changes were observed. For example, we found that a peptide with a more mature appearance, E(Glu→pyro-Glu)NRNFLRFamide, had a higher abundance in the fed state at 0 h (p-value ≤ 0.05; **Fig. S2A**). Another seemingly mature peptide, Q(Gln→pyro-Glu)GNFLRFamide, displayed a decreased fold change at 30 min (**Fig. S2B**). Interestingly, the 1 h time point revealed no significant changes among the isoforms (**Fig. S2C**), whereas the relative abundance changes were variable across the latter three time points and isoforms (**Fig. S2D-F**). Though less dramatic, differences among isoforms were also present in the TG (**Fig. S3**), largely occurring in modified forms of the peptide. By contrast, the fold changes in the STNS (**Fig. S4**) and SG (**Fig. S5**) were more subtle. Altogether, these findings highlight the subtle differences that can give rise to substantial changes in neuropeptide signaling, specifically in response to feeding (**Supplementary File 2**).

Across the SG, STNS, TG, and PO, a feeding state–dependent change in expression of SPYAFGLamide was also observed (**Fig. 3B**). In each tissue, the general trend revealed an upregulation of peptide expression immediately following feeding, followed by a downregulation after the 1 h time point. This pattern suggests a peptide that acts rapidly but then quickly returns toward baseline. A similar trend is evident for KMYDFGLamide, although the return to baseline is slightly slower in the TG (**Fig. 3C**). Both peptides bare the FGLamide sequence, consistent with allatostatin A peptides, and perhaps represent a conserved component of the feeding process.^37^ Finally, GIYGFGL, also containing the FGLamide motif, displayed significant changes exclusively within the STNS. Given the central role of the STNS in feeding-related motor patterns, this suggests that GIYGFGL may be more locally acting (**Fig. 3D**). This peptide is sparsely documented elsewhere, suggested to be an incomplete penaeustatin peptide, only previously identified in the tiger prawn (*Penaus monodon*).^38^ Once more, this highlights the highly conserved functionality that is deeply dynamic and tuned by subtle sequence deviations.

To probe this further we evaluated abundance differences among three sets of peptides that differed by a single residue. Focusing on allatostatin A peptides, we noted that SPY**A**FGL and SPY**E**FGL differ only at the central amino acid, yet this modest change was associated with distinct expression patterns at 0 min and 1 h post-feeding: the E-variant was upregulated immediately post-feeding, whereas the A-variant became upregulated at 1 h (**Fig. 4A**). To determine if this impact would also be apparent with a modified peptide, we compared Q(Gln→pyro-Glu)R**A**YSFGLamide and Q(Gln→pyro-Glu)R**P**YSFGLamide, which differ at the third amino acid residue. These isoforms displayed very similar expression profiles, with the exception of the 6 h time point (**Fig. 4B**). At 6 h, Q(Gln→pyro-Glu)R**A**YSFGLamide was upregulated, while its P-variant counterpart was downregulated, suggesting more sustained activity that coincides with foregut clearance.^5^ Additionally, a dramatic shift in expression between E(Glu→pyro-Glu)PY**A**FGL and E(Glu→pyro-Glu)PY**E**FGL was also observed at 30 min post-feeding (**Fig. 4C**). Collectively, these examples emphasize that even single-residue substitutions can markedly reshape peptidomic responses. This conclusion is further supported by previous work illustrating differential isoform expression within the PO.^3^

**Figure 4.**
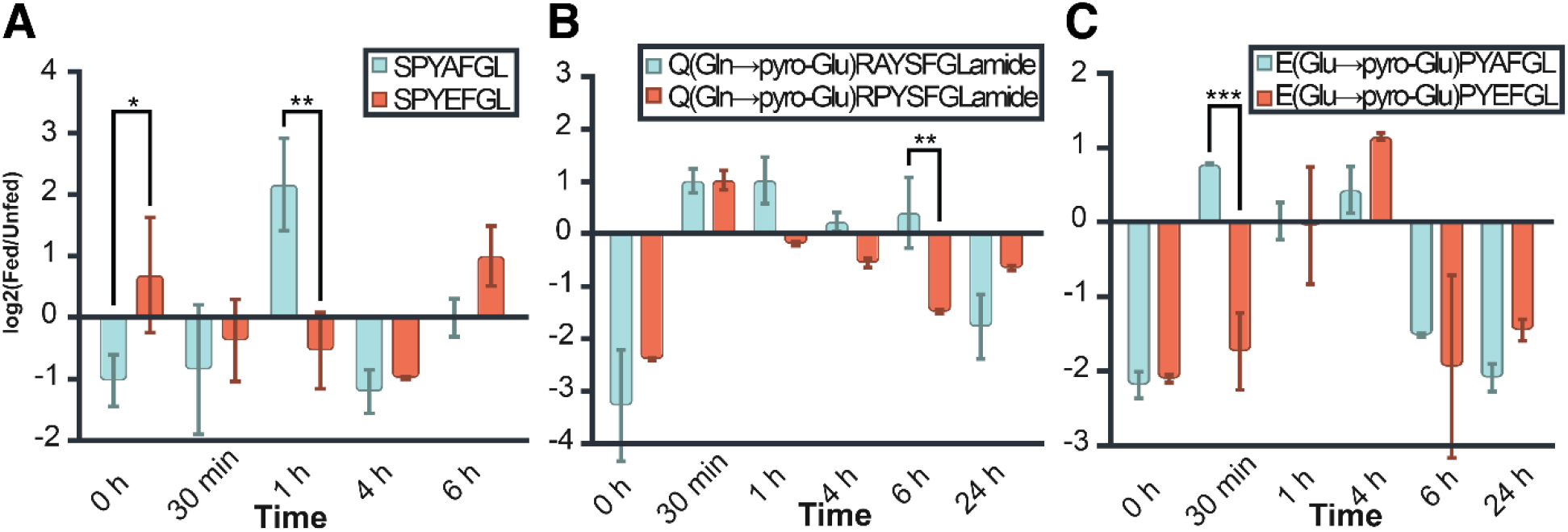
The temporal expression of peptide isoforms within the pericardial organ (PO), where peptides differ by a single amino acid. **A)** SPYAFGL and SPYEFGL, **B)** Q(Gln→pyro-Glu)RAYSFGLamide and Q(Gln→pyro-Glu)RPYSFGLamide, and **C)** E(Glu→pyro-Glu)PYAFGL and E(Glu→pyro-Glu)PYEFGL. Bars represent the average of three technical replicates ± standard deviation. Significance was evaluated via two-way ANOVA with Bonferroni’s multiple comparison test.

Honing in on the influence of modifications, we also evaluated the amidated and non-amidated forms of two peptides. Interestingly, for GHYNFGL (Allatostatin-A; **Fig. S6A**) and FVGGSRY (RYamide; **Fig. S6B**), the overall temporal trends were retained between modified and unmodified forms; however, the amidated form revealed less dramatic shifts in regulation, likely indicative of a tightly regulated system.

To appreciate higher level shifts in up- and down-regulation, we first turned to the PO, which revealed a clear temporal cascade: peptides were largely upregulated within the first hour post-feeding (**Figs. 5A-C**) and largely switching to a downregulated state by 4 h post-feeding (**Fig. 5D**). By 24 h, peptide levels were predominantly downregulated, consistent with the return to a basal physiological state (**Fig. 5E**). Digging further, we evaluated a peptide with highly significant changes during the first three timepoints, as these appeared to be a critical timepoint of peptide up-regulation, peptide FVGGSRYamide (RYamide; **Fig. 5F**). Notably, the peptide appears to have a lower intensity in the fed state at 0 h, shows marked increases at the 30 min and 4 h, with no significant changes at 6 h and 24 h post-feeding. This pattern suggests that FVGGSRYamide is rapidly mobilized and then tightly suppressed as active feeding-related processes resolve. This is further supported with the finding that this peptide has previously been observed in environmental perturbation studies in response to temperature and salinity stress, perhaps implying a homeostatic function.^25, 26^

**Figure 5.**
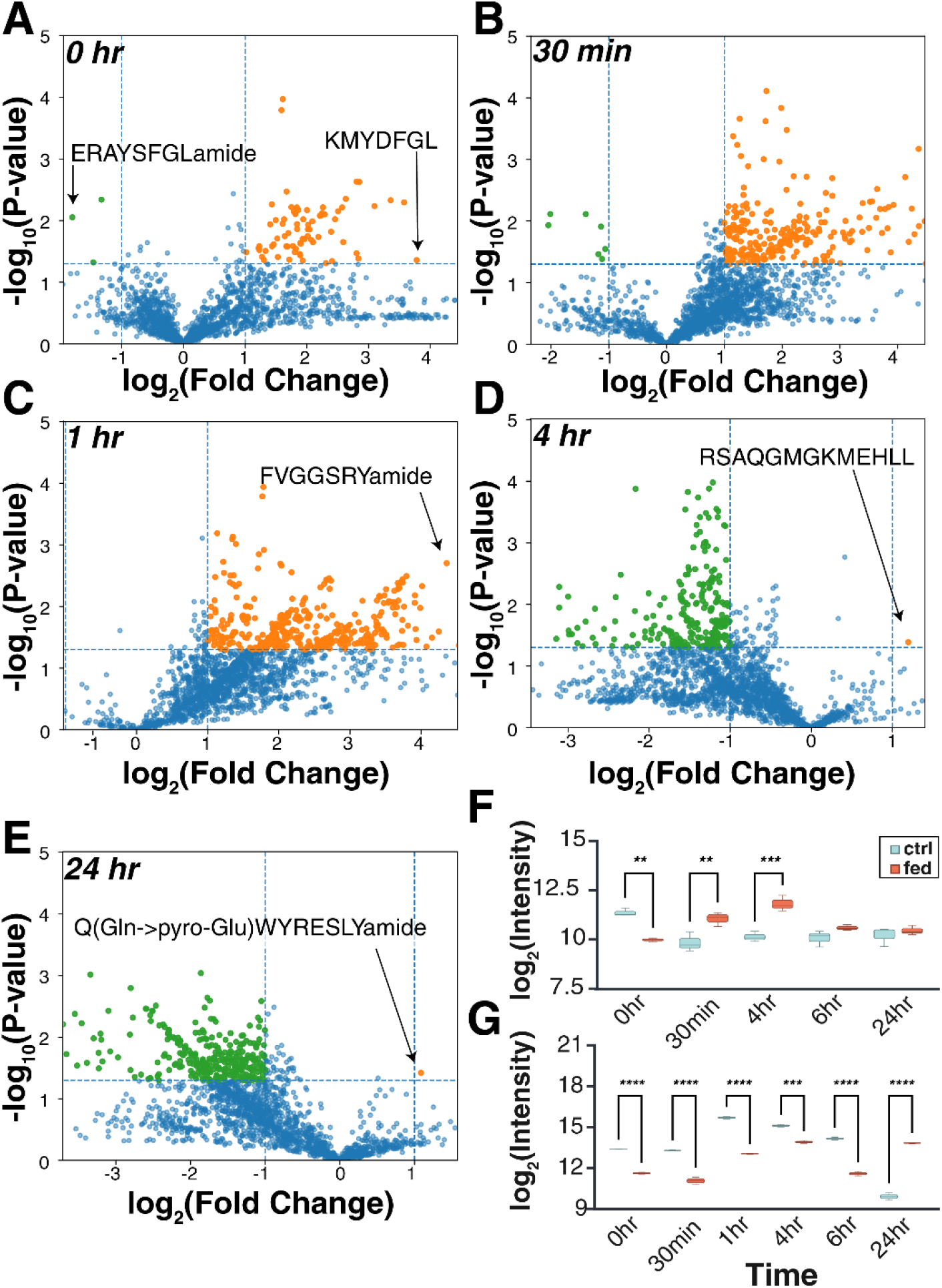
Global analysis of up (orange) and down-regulated (green) peptide expression in the PO at **A)** 0 h, **B)** 30 min, **C)** 1 h, **D)** 4 h, and **E)** 24 h post-feeding. **F)** Expression of FVGGSRYamide and **G)** SDRNYLRF with respect to time. No significantly down-regulated peptides were identified at 1 hr post-feeding.

Additionally, peptide SDRNYLRF (RFamide) was the only peptide showing significant differences between fed and unfed states at every time point (**Fig. 5G**). Interestingly, SDRNYLRF was consistently downregulated across all early and mid-time points but showed a dramatic upregulation by 24 h. While the general expectation is that hormonal signaling should return to baseline by this stage, this observation suggests a prolonged regulatory window. This can point to particular responses, such as extended neuromodulatory activity, a delayed compensatory response, or even additional physiological roles which emerge only after digestion is complete. While SDRNYLRF has previously been identified, its functionality has not previously been investigated.^1, 35^

Previously, PO changes with response to feeding were observed in *Homarus americanus*,^35^ *Cancer borealis*,^1^ *Callinectes sapidus*,^3^ *Carcinus maenas*,^3^ and others. Specifically, neuropeptides are found to be released from the PO into the hemolymph, and subsequently sent through the stomatogastric ganglion (STG) to the brain.^25, 39^ With the TG and PO centralized around cardiac function, it was initially hypothesized that their neuropeptide expressions with respect to feeding would be similar. Thus, it was extremely intriguing that the PO and TG had virtually opposite intensities at each time point (**Fig. S7**). The TG, contributing to the *C. borealis* central nervous system (CNS), seems to exhibit distinct activity from the PO, which engages with the STG. ^25^ More likely, it appears that the TG displays feedback loops with the brain, another component of the CNS.

While the observation of peptidomic trends in the PO and TG were pronounced, the STNS (**Fig. S8**), brain (**Fig. S9**), and SGs (**Fig. S10**) proved much more discrete. In prior studies, we have shown that integration of a “boosting” channel, effectively front-loading peptides into a specific channel, can enhance identification rates by enhancing low-abundance signals.^40^ Although not explicitly used here, such boosting may have occurred implicitly due to the TG’s significantly larger tissue mass, potentially increasing TG detection sensitivity relative to other tissues.

To further probe this phenomenon, we examined proctolin, a peptide that is ubiquitous within neuropeptidomics, notoriously difficult to detect due to its extreme susceptibility to degradation.^41, 42^ The active form of this peptide, RYLPT, lacks PTMs and stimulates foregut musculature, making its regulation highly relevant to feeding behaviors.^1, 43-45^ Elegant studies have been performed to probe the chemical perturbations of this peptide, though the characterization of modified forms, to our knowledge, has not been completed.^42^ One study in *Drosophila* hypothesized that the precursor of proctolin was capable of undergoing amidation, but no such studies have been conducted in crustacea.^45^ Thus, much work must be employed to evaluate if there is a possibility for the amidated form of proctolin to be bioactive.

Interestingly, we observed no significant changes in the unmodified variant of RYLPT within the SGs, which was not detected in any other tissue (**Fig. S11A**). This was particularly interesting that, previously, this peptide was found to have no significant changes in abundance.^1^ This is relevant as, when comparing side-by-side, the isoform RYLPTamide was found in the SGs, with a significant change from the control at 4 h through 24 h post feeding (**Fig. S11B**). Additionally, the unmodified form was not observed in the brain, yet the modified variant was extensively profiled, revealing a significance in change at 1 h, with the fed variant having elevated levels of proctolin (**Fig. S11C**).

We then turned to evaluation of potential inter-tissue crosstalk. By performing principal component analysis, grouping at the tissue level (**Fig. S12A**), distinct clusters could be attributed to the PO and TG, while the brain, SG, and STNS were enmeshed. We further attempted to cluster solely on fed or unfed status, though no clusters could be defined (**Fig. S12B**). When evaluating tissue and fed or unfed status simultaneously, it became apparent that distinct clusters can be assigned within the TG to each fed and unfed state (**Fig. S12C**). This perhaps points to a yet unknown contribution of TG to feeding-related signaling.

By zooming in to determine the peptide-level intersections between tissues (**Fig. S13**), it was evident that peptides AWPHSFS and QRNFLRF (RFamide) were exclusive to the brain. AWPHSFS is a diuretic hormone-44 peptide previously implicated in feeding.^1^ While several peptides were unique to the STNS, no IDs were found to be exclusive just to the brain and STNS, surprising due to their proximity and expected contributions to feeding response. However, in line with prior findings within this study, 51 peptides were shared between the PO and TG. It is also worth noting that 16 peptides were present in all tissues analyzed. Specifically, all but 2 of these peptides were allatostatin A peptides. This was a trend that was also observed in the TG, PO, and SG, where the IDs were largely from allatostatins and RFamides. One interesting note is that many of these IDs were previously implicated in neuromodulation.^38, 46, 47^ These further underscores conserved functionality within the neuropeptidome.

## Conclusions

This work highlights the first multiplexed evaluation of the crustacean neuropeptidome via isobaric labeling. This analytical innovation revealed unexpected findings, including the amplification and improved identification of otherwise elusive peptides. By using this approach to investigate neuropeptide signaling in response to feeding, we define distinct profiles of peptides involved in these processes. We further use this as a testbed to probe the activity of mature peptide isoforms compared to unmodified peptides, as well as the peptide abundance differences of sequences corresponding to the same family, delving as far as peptides differing by just a single amino acid. Finally, we present evidence for a potentially modified expression of the proctolin neuropeptide. Altogether, this illustrates a method capable of being applied to sparse peptide samples from which complex mechanisms can be probed quantitatively.

## Data Availability

All mass spectrometry proteomics data have been deposited to the ProteomeXchange Consortium via the MassIVE partner repository with the dataset identifier: MSV000101048.

## Acknowledgements

This work was supported in part by National Institutes of Health (NIH) through grants R01AG078794 (L.L.), R01DK071801 (L.L.), , and the Research Forward grant by University of Wisconsin - Madison Office of the Vice Chancellor for Research with funding from the Wisconsin Alumni Research Foundation (L.L.). The Orbitrap instruments were purchased through the support of NIH shared instrument grant S10RR029531 and Office of the Vice Chancellor for Research and Graduate Education at the University of Wisconsin-Madison. L.F. was supported in part by the National Institute of General Medical Sciences of the National Institutes of Health under Award Number T32GM008505 (Chemistry–Biology Interface Training Program), the 2024 Eli Lilly and Company/ACS Analytical Graduate Fellowship, and a predoctoral fellowship supported by the NIH, under Ruth L. Kirschstein National Research Service Award (NRSA) from the National Institutes of Health-General Medical Sciences F31GM156104. L.L. would like to acknowledge NIH grants R01AG052324, S10OD028473, and S10OD025084, as well as funding support from a Vilas Distinguished Achievement Professorship and Charles Melbourne Johnson Professorship with funding provided by the Wisconsin Alumni Research Foundation and University of Wisconsin-Madison School of Pharmacy. The authors acknowledge the UWCCC Experimental Animal Pathology Laboratory for their core services, which is supported by P30CA014520. Figures 1, 3, 4, S1, and S11 were generated with BioRender.

## For TOC Only

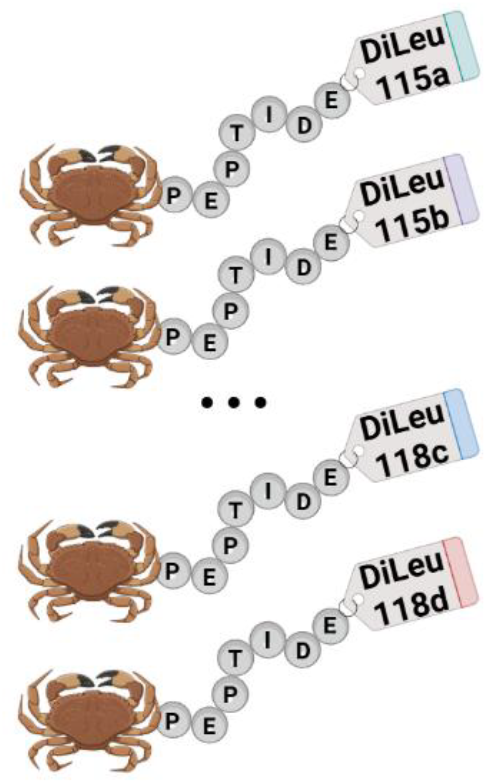

