## Supporting Information for "12-plex DiLeu enables robust quantification of the feeding neuropeptidome"

**Table of Contents**

Supplemental Figures (located within this document)


- **Figure S1:** Comparison of EndoGenius and false discovery rate
- **Figure S2:** Evaluation of NFLRFamide peptides in pericardial organs
- **Figure S3.** Evaluation of NFLRFamide peptides in thoracic ganglion
- **Figure S4:** Evaluation of NFLRFamide peptides in the stomatogastric nervous system
- **Figure S5:** Evaluation of NFLRFamide peptides in the sinus glands
- **Figure S6:** Comparison of amidated and non-amidated peptides
- **Figure S7:** Global peptide analysis in thoracic ganglion
- **Figure S8:** Global peptide analysis in stomatogastric nervous system
- **Figure S9:** Global peptide analysis in the brain
- **Figure S10:** Global peptide analysis in the sinus glands
- **Figure S11:** Evaluation of proctolin neuropeptides
- **Figure S12:** Characterization of peptides via principal-component analysis
- **Figure S13:** Upset plot of all identified neuropeptides


Supplemental Tables (located within this document)


- **Table S1:** Isotopic interference correction table


Supplemental Files


- **Supplemental File 1:** Comparison of peptides identified in all tissues
- **Supplemental File 1:** Intensities of NFLRFamide peptides

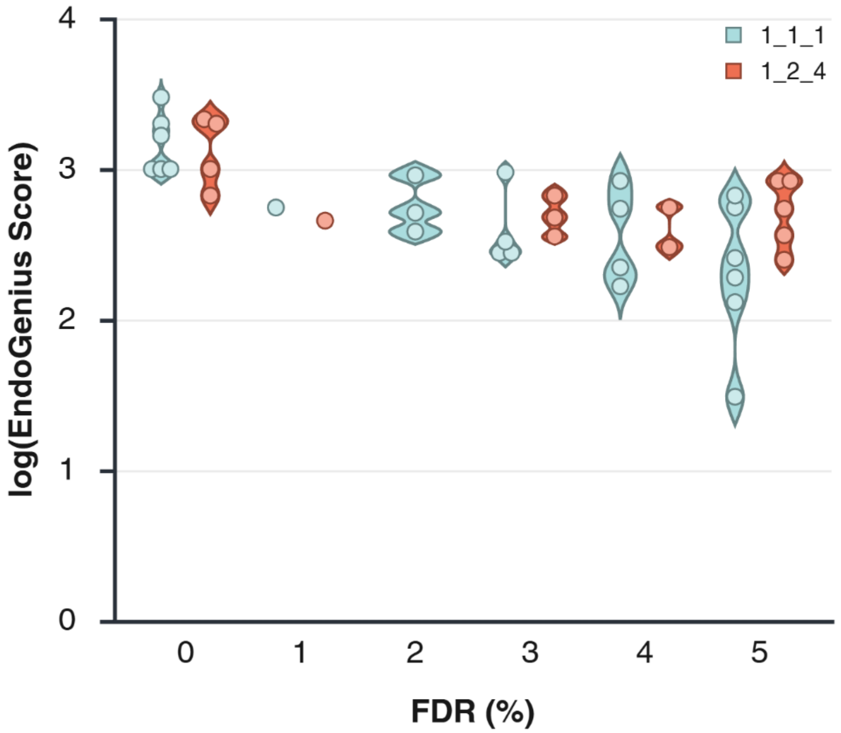


**Figure S1:** Relationship between FDR and the log(EndoGenius Score) with relation to 15 spectral datasets at an equal and variating ratios. Blue datapoints represent samples prepared in an equal ratio across all channels (1:1:1:1:1:1:1:1:1:1:1:1), while red datapoints represent samples prepared in variable ratios across all channels (1:2:6:10:12:12:10:6:4:2:1).

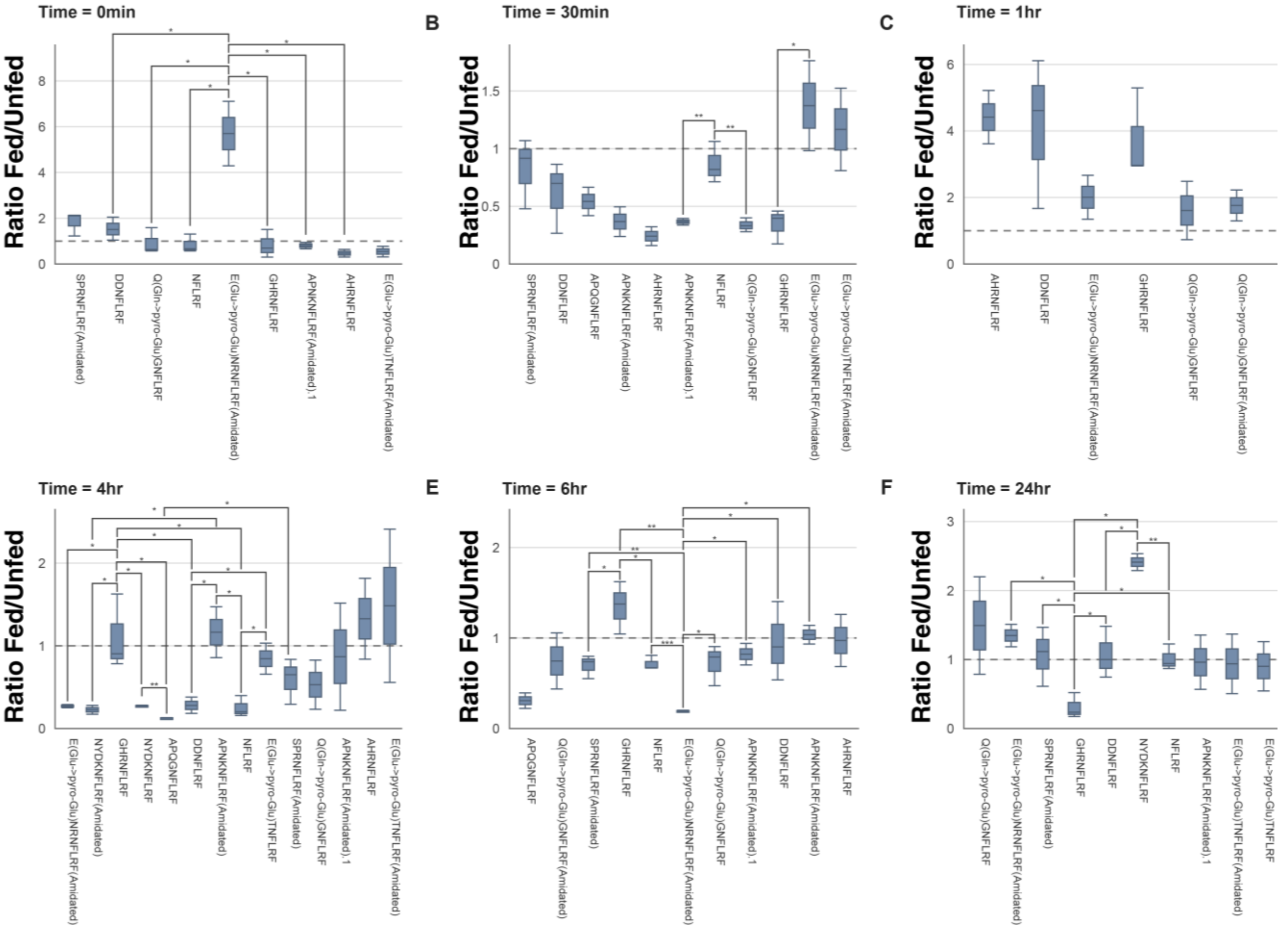


**Figure S2:** Characterization of peptides including the NFLRFamide motif within the POs across **A)** 0 h, **B)** 30 min, **C)** 1 h, **D)** 4 h, **E)** 6 h, and **F)** 24 h time points. Pairwise comparisons between fragments were performed using two-tailed Welch's t-tests.

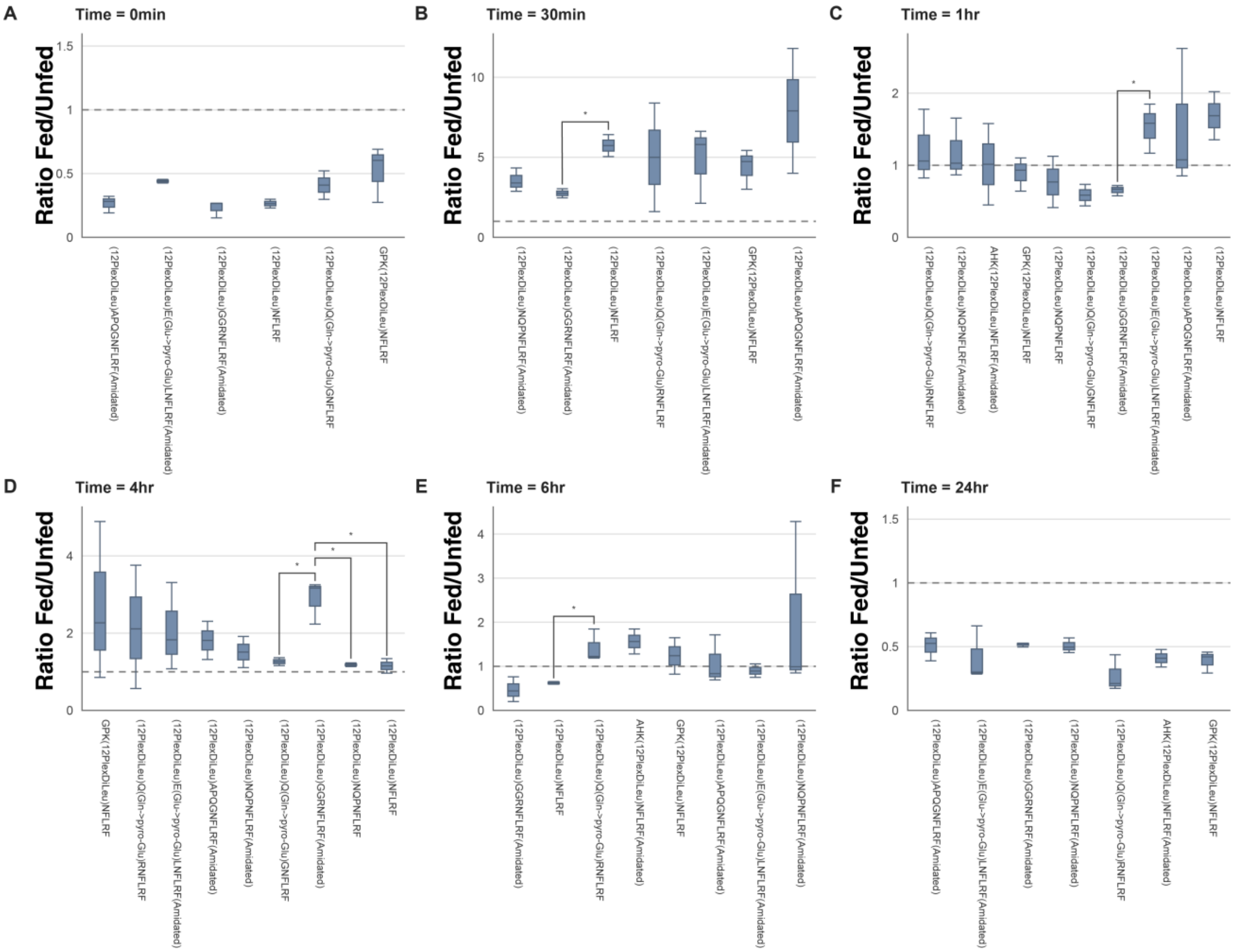


**Figure S3:** Characterization of peptides including the NFLRFamide motif within the TG across **A)** 0 h, **B)** 30 min, **C)** 1 h, **D)** 4 h, **E)** 6 h, and **F)** 24 h time points. Pairwise comparisons between fragments were performed using two-tailed Welch's t-tests.

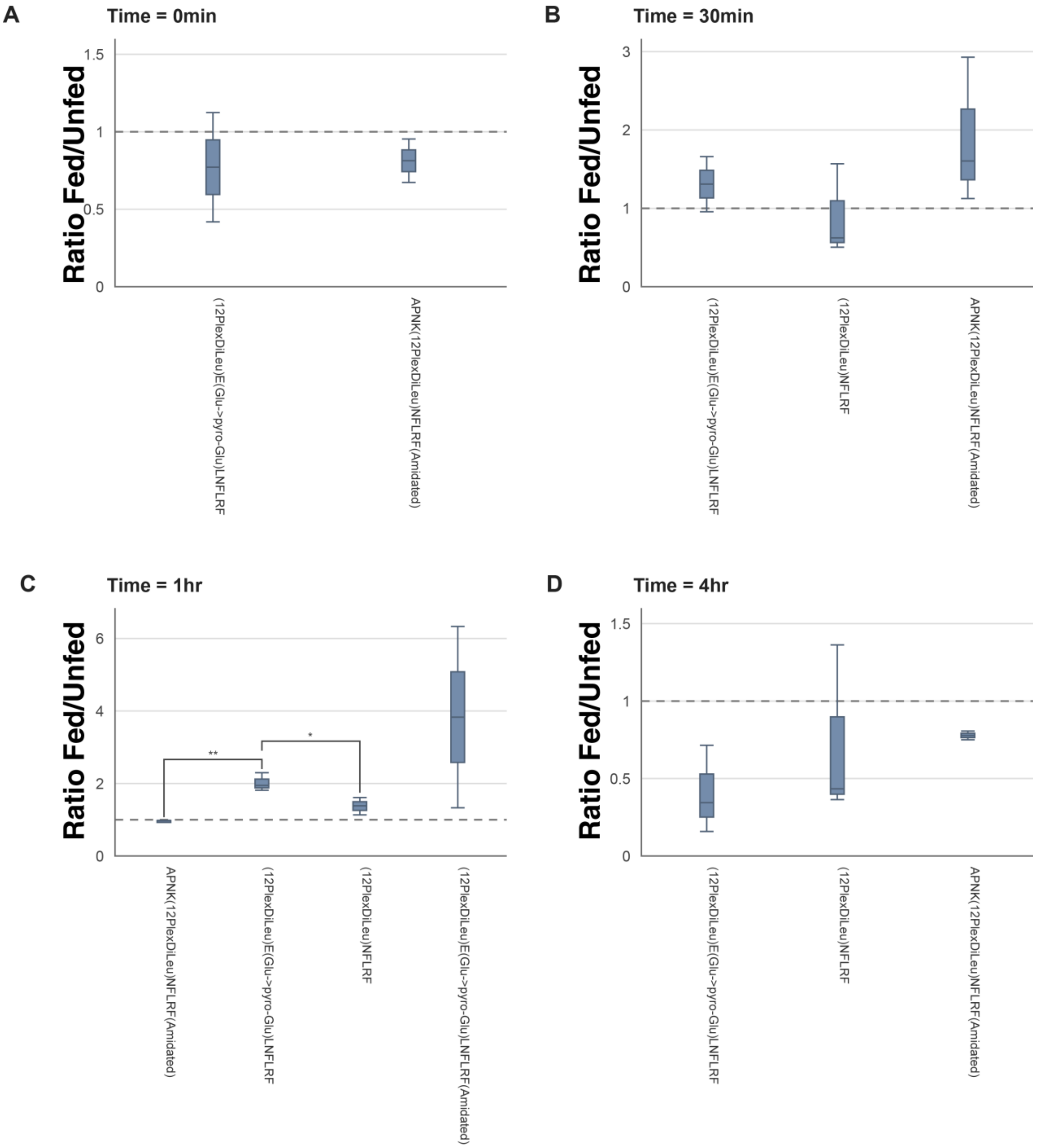


**Figure S4:** Characterization of peptides including the NFLRFamide motif within the STNS across **A)** 0 h, **B)** 30 min, **C)** 1 h, **D)** 4 h points. The 6 h and 24 h time points did not have detected NFLRFamide peptides. Pairwise comparisons between fragments were performed using two-tailed Welch's t-tests.

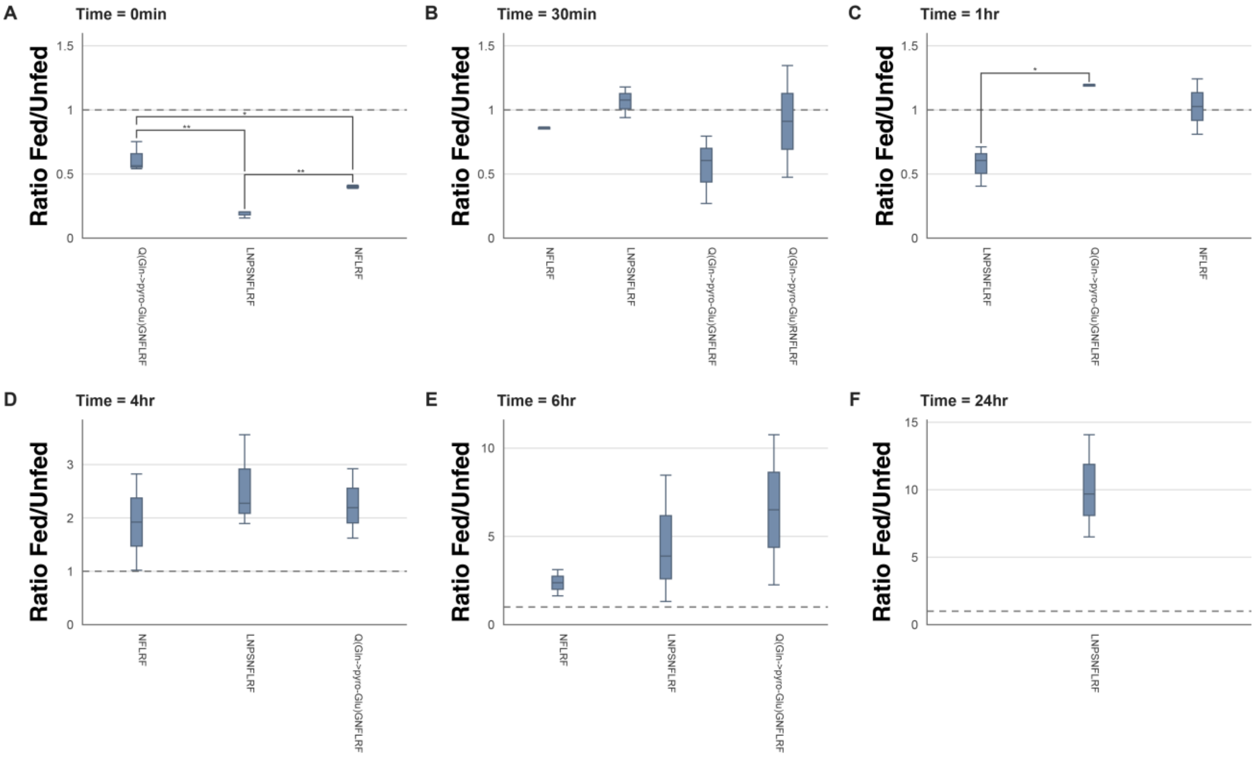


**Figure S5:** Characterization of peptides including the NFLRFamide motif within the SGs across **A)** 0 h, **B)** 30 min, **C)** 1 h, **D)** 4 h, **E)** 6 h, and **F)** 24 h time points. Pairwise comparisons between fragments were performed using two-tailed Welch's t-tests.

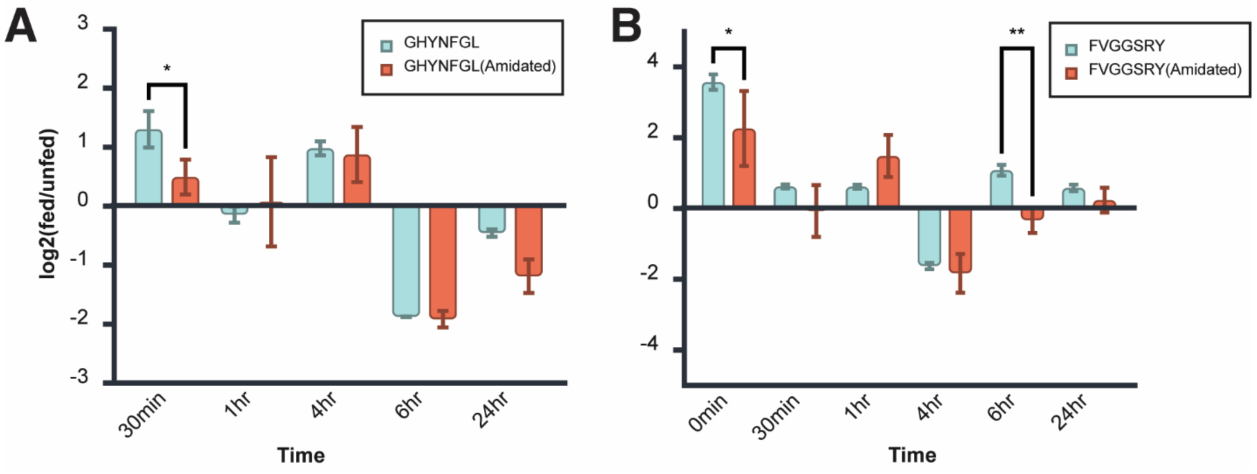


**Figure S6:** Evaluation of amidated (blue) and non-amidated (red) forms **of A)** GHYNFGL and **B)** FVGGSRY across the timepoints.

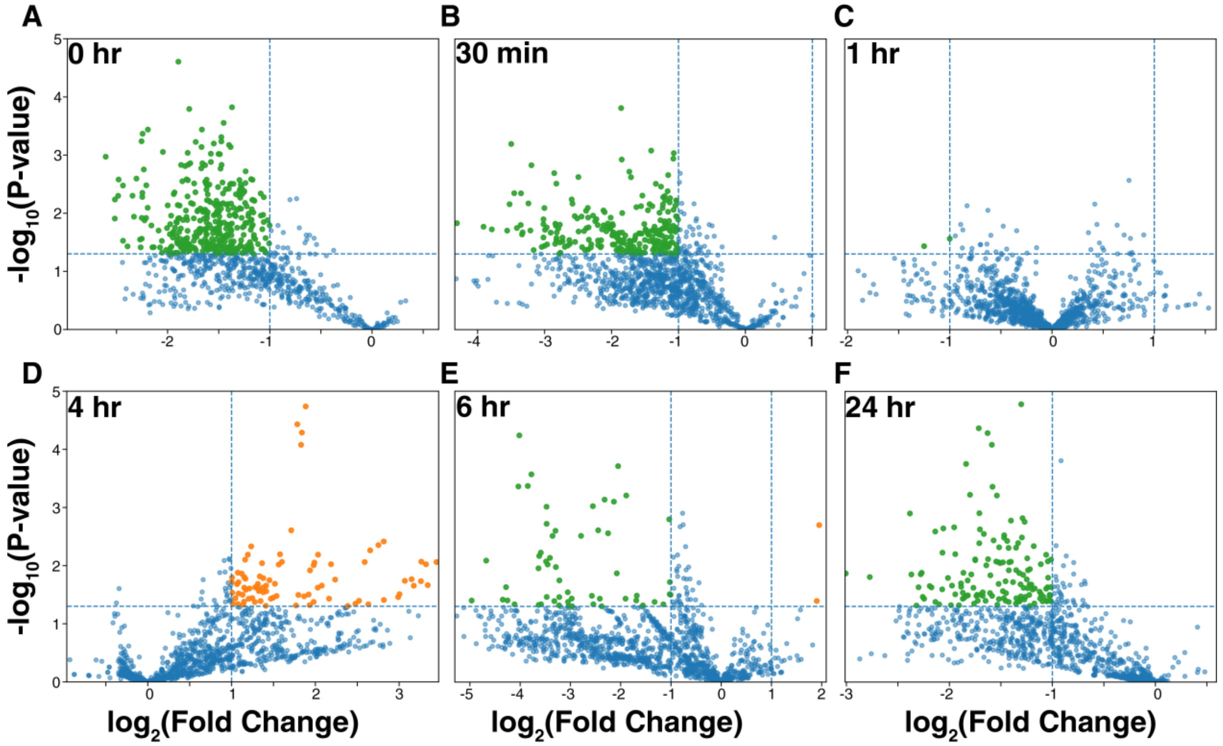


**Figure S7:** Global analysis of up (orange) and down-regulated (green) peptide expression in the TG at **A)** 0 h, **B)** 30 min, **C)** 1 h, **D)** 4 h, **E)** 6 h, and **F)** 24 h post-feeding.

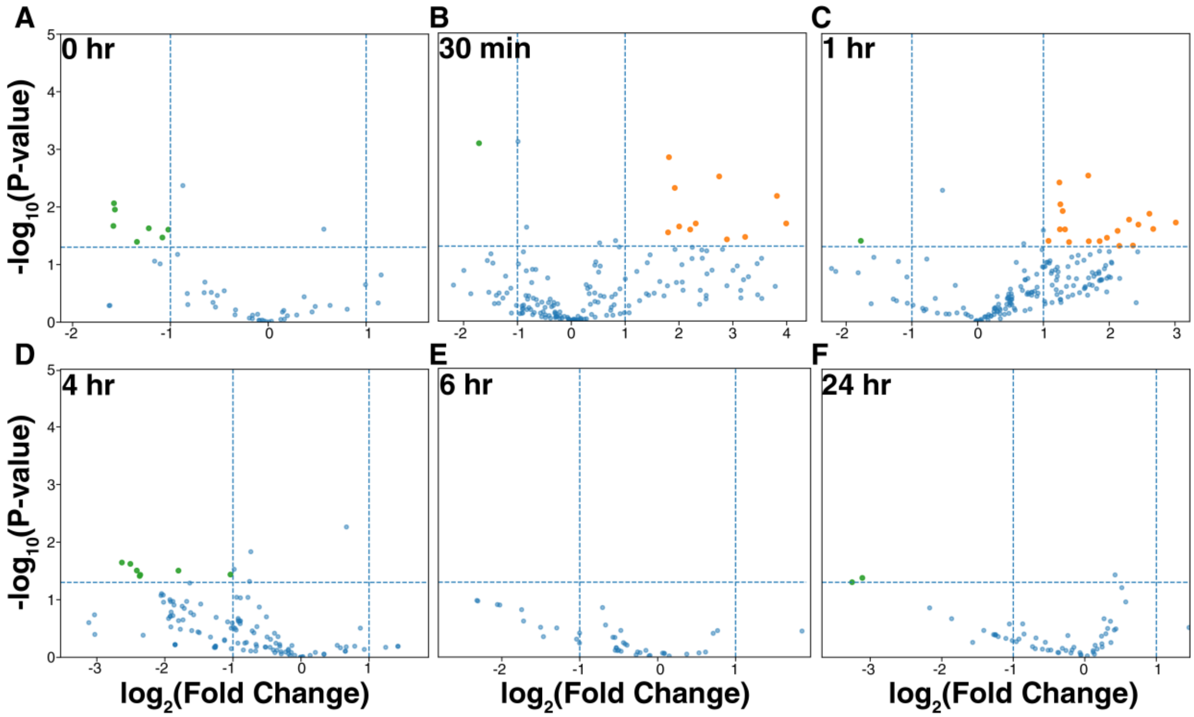

**Figure S8:** Global analysis of up (orange) and down-regulated (green) peptide expression in the STNS at **A)** 0 h, **B)** 30 min, **C)** 1 h **D)** 4 h, **E)** 6 h, and **F)** 24 h post-feeding.

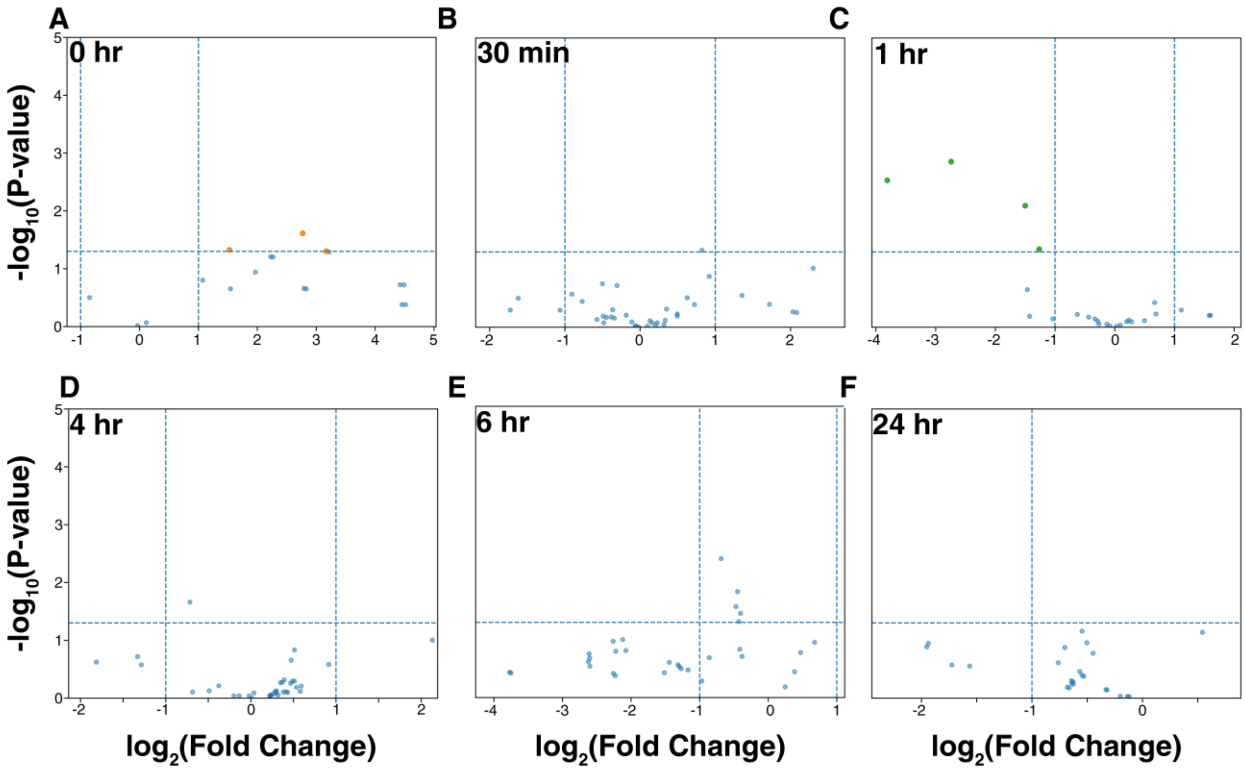


**Figure S9:** Global analysis of up (orange) and down-regulated (green) peptide expression in the brain at **A)** 0 h, **B)** 30 min, **C)** 1 h, **D)** 4 h, **E)** 6 h, and **F)** 24 h post-feeding.

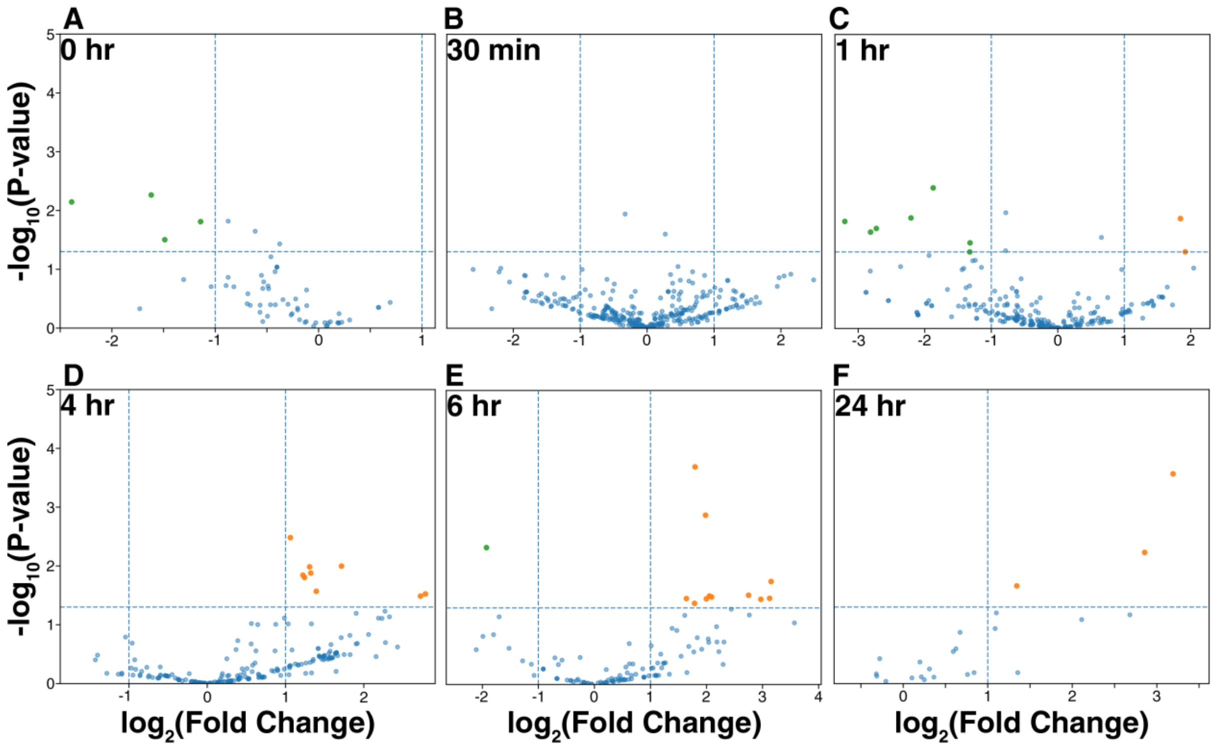


**Figure S10:** Global analysis of up (orange) and down-regulated (green) peptide expression in the SGs at **A)** 0 h, **B)** 30 min, **C)** 1 h, **D)** 4 h, **E)** 6 h, and **F)** 24 h post-feeding.

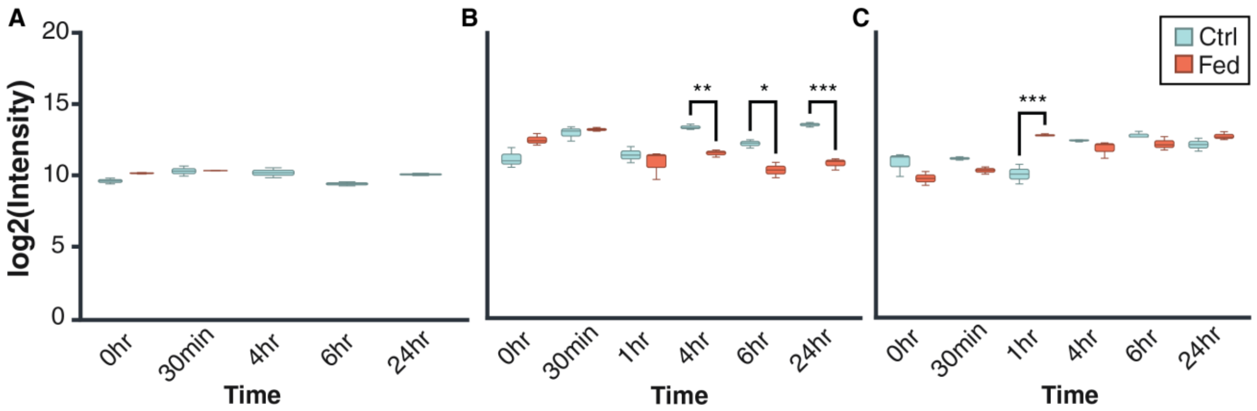


**Figure S11:** Evaluation of proctolin (*i.e.* RYLPT) expression in the SGs in **A)** a non-amidated form and **B)** when amidated. **C)** Expression of amidated proctolin in the brain.

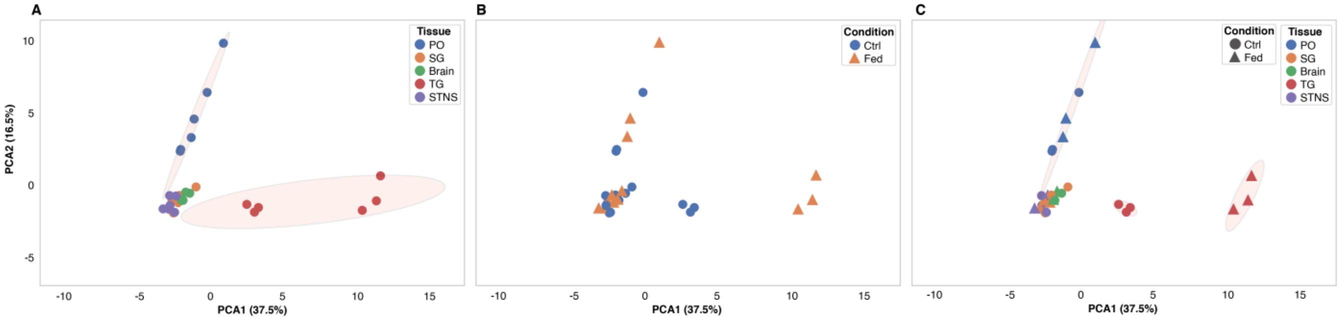


**Figure S12:** Global characterization of peptides via principal component analysis, annotated on the basis of **A)** tissue type, **B)** feeding condition, and **C)** combination of tissue type and feeding condition.

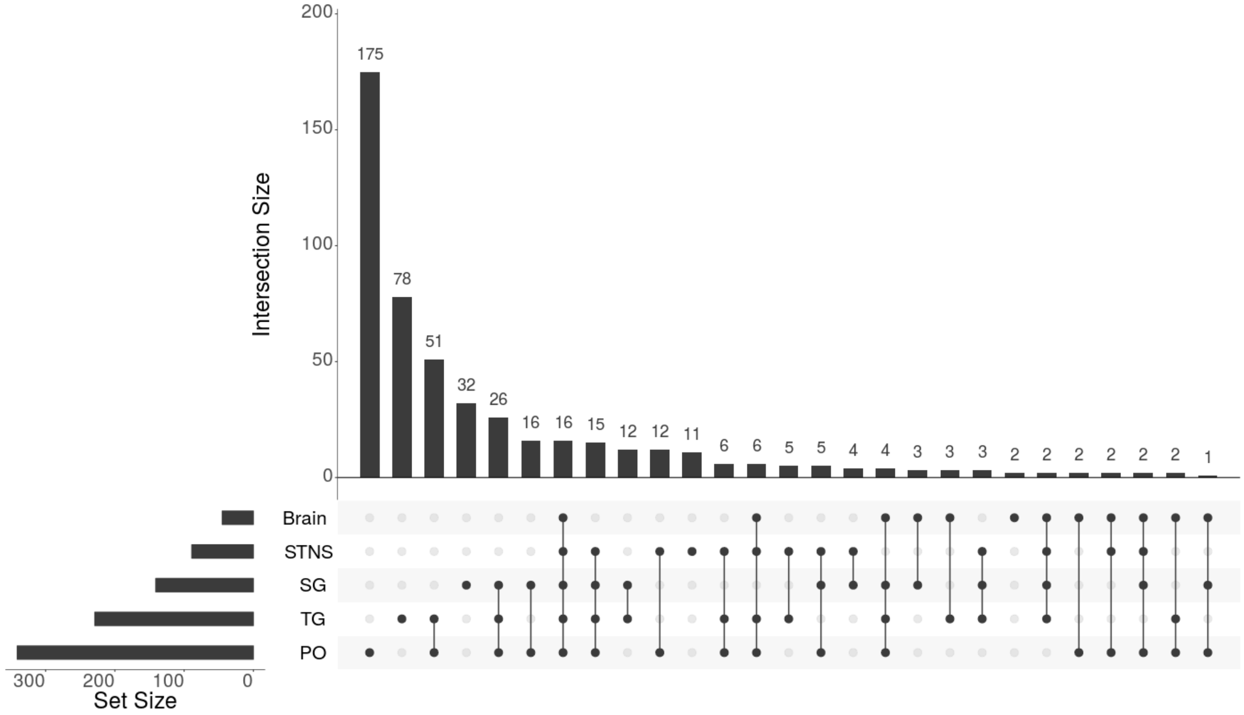


**Figure S13:** Upset plot defining the intersection of peptides defined in all tissues analyzed.

| | 115a | 115b | 116a | 116b | 116c | 117a | 117b | 117c | 118a | 118b | 118c | 118d |
|---|---|---|---|---|---|---|---|---|---|---|---|---|
| **115a** | 100.00% | | 0.12% | | | | | | | | | |
| **115b** | | 100.00% | 0.65% | 2.02% | | | | | | | | |
| **116a** | 7.06% | | 100.00% | | | 1.25% | | | | | | |
| **116b** | | 6.10% | | 100.00% | | 1.90% | | | | | | |
| **116c** | | | | | 100.00% | | 0.46% | 0.97% | | | | |
| **117a** | | | 2.51% | | | 100.00% | | | 2.37% | | | |
| **117b** | | | | | 0.20% | | 100.00% | | | 0.51% | | |
| **117c** | | | | | 3.05% | | | 100.00% | | 0.50% | 1.00% | |
| **118a** | | | | | | 1.67% | | | 100.00% | | | |
| **118b** | | | | | | | 7.35% | | | 100.00% | | |
| **118c** | | | | | | | | 2.60% | | | 100.00% | |
| **118d** | | | | | | | | | | | | 100.00% |

**Table S1:** Isotopic interference imparted by DiLeu tags, utilized for normalization.
